# Excess entropies reveal higher organization levels in developing neuron cultures

**DOI:** 10.1101/2020.03.05.979310

**Authors:** Norbert Stoop, Ralph L. Stoop, Karlis Kanders, Ruedi Stoop

## Abstract

Multi-component systems often exhibit dynamics of a high degree of complexity, rendering it difficult to assess whether a proposed model’s description is adequate. For the multitude of systems that allow for a symbolic encoding, we provide a symbolic-dynamics based entropy measure that quantifies the degree of deviation obtained by a systems’s internal dynamics from random dynamics using identical average symbol probabilities. We apply this measure to several well-studied theoretical models and show its ability to characterize differences in internal dynamics, thus providing a means to accurately compare model and experiment. Data from neuronal cultures on a multi-electrode array chip validate the usefulness of our approach, revealing inadequacies of existing models and providing guidelines for their improvement. We propose our measure to be systematically used to develop future models and simulations.

## INTRODUCTION

The assessment of whether the dynamics of a system composed of many components is correctly described by a model, is facilitated if the dynamics is of an inherently simple nature. The latter may be the case if specific simplifying features, such as criticality [1], synchronization [2, 3], or corresponding architectural constraints [4] are present. It has been argued that biological neural networks host such beneficial properties [5–8], but whenever this fails to dominate the behavior [9], such an assessment turns into a difficult task. Choosing neural networks as the showcase, we demonstrate how symbolic dynamics-founded ‘excess entropies’ 1) measure how much simulations differ from the target processes, 2) uncover model inadequacies and 3) provide guidelines for model set-up and improvement. This opens a new gateway for the analysis and description of complex dynamical processes that the traditionally used measures do not offer.

## EXCESS ENTROPY ANALYSIS

Formal languages, first used to classify computer and human spoken languages [10–13], have recently added important insight in dynamical systems, in theory and in applications [14–17]. Dynamical systems map a generically chosen initial condition into a sequence **S** = {*s*(*t*_1_), *s*(*t*_2_), …} of a countable number of observational states labeled by symbols *s* ∈ *S* that the system resides in at a given time *t*_*j*_. In the neuronal culture experiments that later we will use to validate our measure, the role of the observational states is played by activated multi-electrode array (=MEA) electrodes (we will therefore use ‘symbols’ and ‘electrodes’ synonymously). **S** can be seen as generated according to a set of rules imposed by a grammar from one of the Chomsky language classes t-0–t-3 [10], ordered by the ease of capturing complex relations among the language elements. Alternatively, the complexity of the structure of **S** can be measured by the difficulty of the prediction of future system states from past states [16], and the amount of computation performed by this process can be measured in objective terms [18]. If each state is visited at random with probabilities **p** = {*p*_*j*_}, 1 ≤ *j* ≤ *s*, the probability for observing a sequence of symbols **S** of length *N* (and any permutation thereof) is given by the multinomial distribution

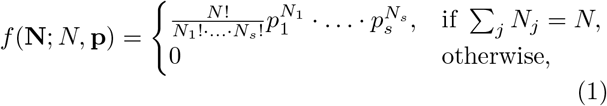

where the random variable vector **N** = {*N*_*j*_} denotes the number of observations for each state *j* and where the true state probabilities are approximated by the observed sample probabilities **p** = **N**/*N*. *f*(**N**; *N*, **p**) does not embrace the order by which the states are visited and therefore does not retain information regarding the underlying system dynamics. To capture the latter aspect, we consider at each position *n* of the sequence the *walk-through probability* [17] which evaluates for each symbol *s*_*j*_ (*n*) the probability that a random walk starting with symbol *s*_1_ will pass through symbol *s*_*j*_ (i.e. *P*_*in*_(*s*_*j*_)), times the probability that a random walk starting at *s*_*j*_ will end with symbol *x*_*L*_ (ie, *P*_*out*_(*s*_*j*_))

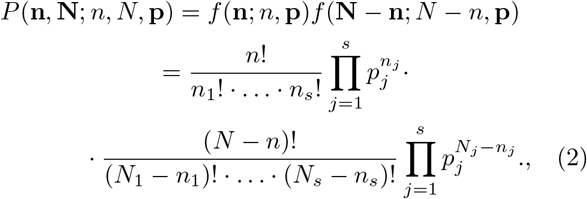

where **n** = {*n*_*j*_} denotes the number of times each state is observed up to position *n*, 0 ≤ *n* ≤ *N*. Using the generalized Vandermonde identity for multinomials [19],

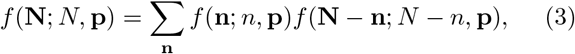

where the sum is restricted to **n** such that ∑_*j*_ *n*_*j*_ = *n*, we obtain the conditional

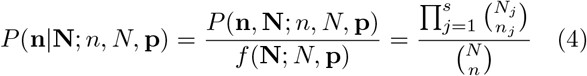

measuring the probability to observe symbol counts **n** in a subsequence of length *n*, given a sequence with total symbol counts **N**. For ease of notation, we will abbreviate in the following *P*(**n|N**; *n*, *N*, **p**) as *P*(**n|N**); *P*(**n|N**) is the multivariate hypergeometric probability distribution (i.e., for sampling without replacement) with expectation value 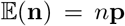. Moreover, *P*(**0|N**) = *P*(**N|N**) = 1, due to symmetry, and *P*(**n|N**) < 1 for all 0 < *n* < *N*, as for any substring of length 0 < *n* < *N*, *P*(**n|N**) contains some information about the actual permutation of symbols in the sequence, whereas *P*(**0|N**) does not.

Introducing the *walk–through entropy h*(*n*) by

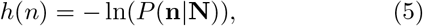

the maximum of *P*(**n|N**) attained for 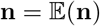 provides a lower bound for *h*(*n*), realized by any sequence that comprises, at any intermediate position *n*, the observed states 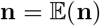 (for details see our Supplemental Material SM). As illustration, consider a sequence of length *N* consisting of three states 1, 2, and 3 with equal probabilities *p*_1,2,3_ = 1/3. *h*(*n*) is minimal if every state occurred precisely *n*/3 times in a subsequence of length *n*, i.e. if the states are visited periodically with the highest possible frequency (Fig. 1a, blue). In contrast, *h*(*n*) will be largest for the lowest frequency sequence where state 1 is exclusively visited in the first third of the sequence, followed by states 2 and 3 in the remaining two thirds (Fig. 1a, red). Even after interchanging 2*N* randomly chosen pairs of states in either sequence, *h*(*n*) still effectively discriminates between the low and high frequency sequences (Fig. 1b, red vs. blue). In particular, for random walks on the three states 1, 2, 3, the expected value 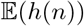 is larger than the lower bound (Fig. 1b, black vs. green).

**FIG. 1.**
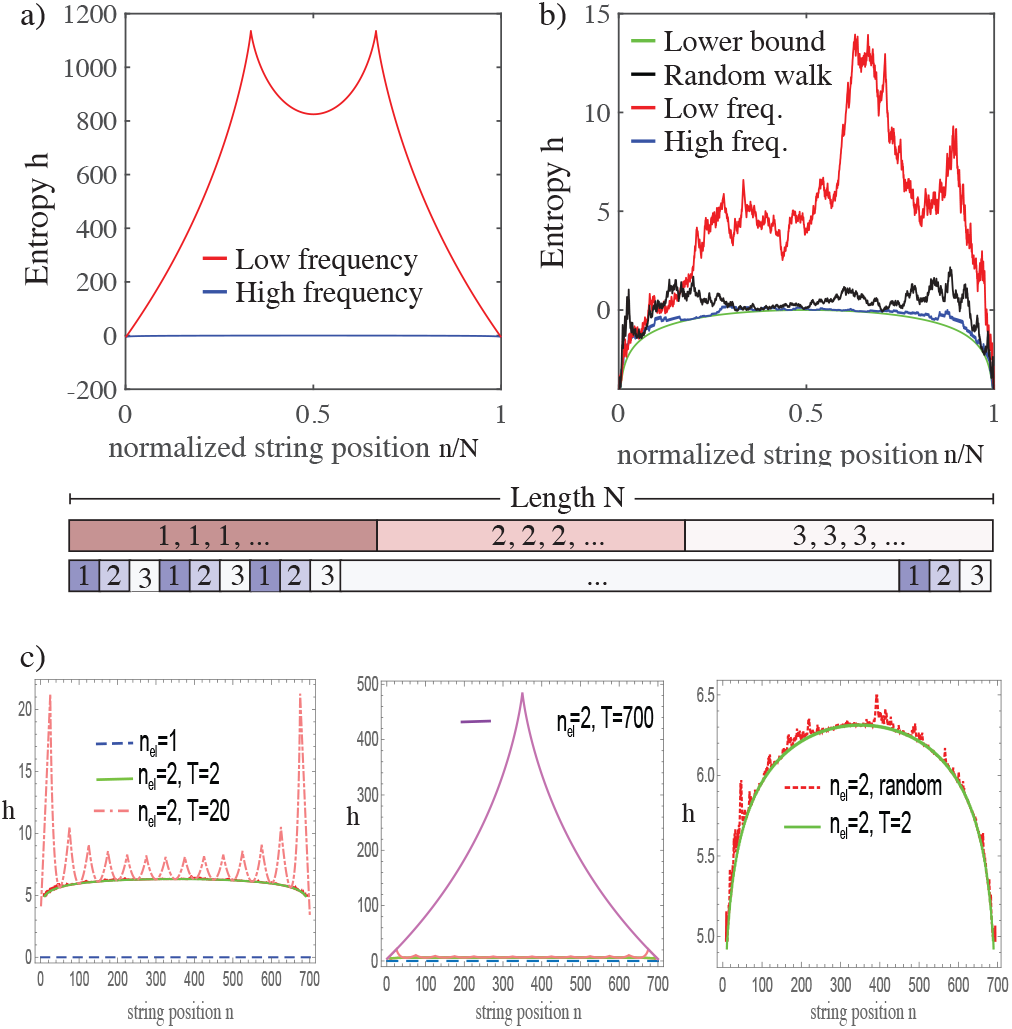
Walk-through entropy *h* evaluated along symbolic strings. a) Low and high frequency compositions of symbols {1, 2, 3}. b) Randomly perturbed low (red) and high (blue) frequency sequences of a) compared to a true random walk (black) and the lower bound (green), cf. SM. c) Artificial strings of length 700 from *n*_*el*_ states and periodicity *T*. Left: Single state *ω* = {*a*, *a*, *a*,…} (*n*_*el*_ = 1, blue dashed line) compared to *n*_*el*_ = 2 active electrodes of periodicity 2 (*T* = 2, *ω* = {*a*, *b*, *a*, *b*, …}, green solid line) and of periodicity 20 (*T* = 20, orange dashed line), and (middle panel) to periodicity 700 (*n*_*el*_ = 2, *T* = 700, violet solid line). Right: Two-state random walk, red dashed line, compared to periodicity 2 (*n*_*el*_ = 2, *T* = 2, green solid line).

*The random-walk baseline* 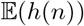 *is thus an insightful means to discriminate between different degrees and qualities of complex behavior: Values of h*(*n*) *below* 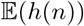 *demonstrate that states are visited in a too orderly fashion, compared to what we expect from a random process; values above* 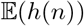 *correspond to sequences with long-wavelength regularity. This not only provides information on how close to given experimental data a simulation is (and whether, upon changing model properties, the modeling is improved); on its own, the level above the random-walk baseline indicates how much of a higher level of ‘ordering’ is present in the data (that could or should be included in the model).*

In Fig. 1c we illustrate these observations by three additional firing patterns. The trivial case with only one symbol (*n*_*e*_ = 1, *ω* = {1, 1, 1,, 1,…}) leads to zero entropy (left subpanel, blue dashed line). Two symbols with each 350 spiking events in a perfect periodic sequence with smallest periodicity *ω* = {1, 2, 1, 2, 1, 2, …} yield, with their uniform distribution of spiking events, a first elevated entropy level (left subpanel: green line solid line). Increasing the periodicity to *ω* = {1, 1, 1, 2, 2, 2, 1, 1, 1, 1, 2, 2, 2,…} leads to alternating piecewise increasing and decreasing regions (left subpanel: orange dashed line for a periodicity of 20), where extrema highlight symbol switching positions, from 1 to 2 or vice versa. The extreme case *ω* = {1, 1, 1,, ….1,, 2, 2,.., 2} leads to a pyramid-shaped *h*-function (middle subpanel, violet solid line). Because this sequence is maximally unlikely, the overall entropy is maximal for this example. Finally, a random walk on two states results in a shape very similar to that of the 2-periodicity sequence (right subpanel: red solid line), where only very small regions of locally increased entropy emerge, due to the randomness of the underlying string generation. These properties suggest to compare the excess entropy obtained from strings from dynamical systems of interest with the excess entropy from corresponding random walks.

Even if all other parameters are held constant, the walk-through probability *P*(**n|N**) depends on the sequence length *N* as well as on the symbol probabilities *p*_*s*_ (cf. SM). To render results comparable across different dynamical processes, we introduce the *excess entropy*

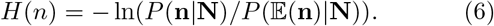

We note that for greatly differing symbol probabilities, this expression still has a - generally weak - asymptotic dependence on *N*. For random processes, we have

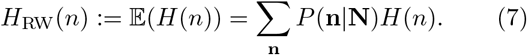

Below we will show how excess entropies detect and distinguish the presence of fine-grained ingredients of random and non-random nature, in real-world vs. simulated data.

## ITERATED MAPS

Because one-dimensional iterated maps reflect many salient properties of complex systems, we first evaluate the excess entropy for the symbolic dynamics of two complementary simple examples. The logistic map [20]

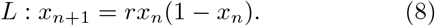

is the key example for how the increase of the nonlinearity leads generically to a universal period-doubling cascade. Starting after *r* = 3 with a period-2, cf. Fig. 2a, the dynamics becomes chaotic at *r*_*_ ≳ 3.56995. To assign a symbolic dynamics, we split the unit interval at *x* = 0.5 into a left and right interval, tagged by symbols 0 and 1, respectively. With the increase of *r*, the period doubling transitions are expressed in cycles of longer periodicity in the symbols 0 and 1 (cf. the back-iteration of the value *x* = 0.5). This is reflected in the measured excess entropy: For *r* < *r*_*_, the excess entropy is essentially 0 (since each symbol appears periodically), but then increases significantly for *r* > *r*_*_ (cf. the two examples of Fig. 2b). The critical role of *r* becomes evident if we consider the mean excess entropy 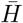 averaged over *N*_*S*_ samples and sequence length *N*,

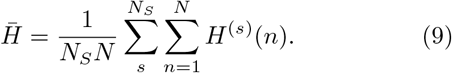

**FIG. 2.**
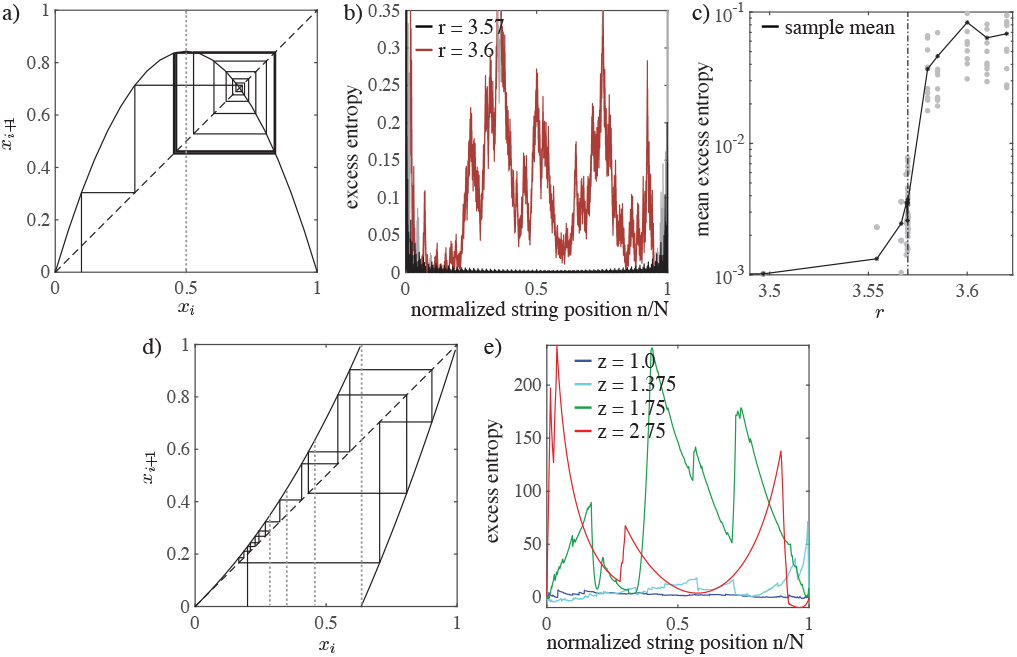
a)-c): Excess entropy of the logistic map. a) Trajectories from a generic initial condition *x*_0_ towards a stable period-2 at *r* = 3.37475. As *r* is increased, at the period-doubling transition to chaos (*r*_*_ ≳ 3.56995), the excess entropy rises sharply (b, c). d)-e) Excess entropy for the intermittent Manneville-Pomeau map. Trajectory for parameter *z* = 2.2, *x*_0_ = 0.2 (d). Dotted vertical lines indicate the interval partition used to associate *x*_*i*_ to discrete states. Large and regularly developing excess entropies are the consequence of the laminar motion induced by the marginal fixedpoint (e).

As may be expected, at *r* ≃ *r*_*_ a sharp increase in 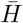 marks the critical transition (Fig. 2c).The excess entropies are, however, much lower than those obtained from random walks *H*_RW_(*n*) (Eq. 7): The sequence mean 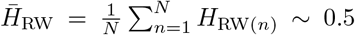, is considerably larger than the mean excess entropy even of the chaotic regime. *Since H is a measure of the imbalance between symbol occurrences, this indicates that even though the dynamics is chaotic, both symbols* 0 *and* 1 *still appear in a fairly balanced fashion; in any subsequence, the probabilities to observe either symbol essentially agree with the overall symbol probabilities*.

As a consequence, we expect *H*(*n*) > *H*_RW_(*n*) for intermittent Pomeau-Manneville dynamics [21]

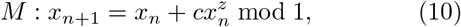

where we choose *c* = 1 (Fig. 2d). For *z* = 1, *M* reduces to the well-known Bernoulli shift [22]. For *z* > 1, it can be understood as a nonlinear extension of the latter, where the behavior in the right-half of the unit interval is Bernoulli-map chaotic; the dynamics in the left-half, however, is dominated by the stability properties of the fixed point *x* = 0. For the symbolic dynamics of *M*, we again split the unit interval guided by the two branches of the map, which provides a natural partition [23]; for *z* > 2 the natural measure becomes non-normalizable. For *z* = 1 (i.e. the Bernoulli-shift), *H*(*n*) is non-zero, but relatively small (Fig. 2e). As *z* increases, *subsequences of intermittent dynamics lead to large values of H*(*n*), *followed by a strong decrease once the dynamics escapes the vicinity of the fixed point x* = 0. Averaging over 10 samples, we find *H*(*n*) ≫ *H*_RW_(*n*), which holds with no apparent dependence on the exponent *z* > 1. The influence of *z* becomes apparent only by the flip ratio *f* defined as the number of sign changes in the increments of *H*(*n*), essentially counting the number of a sequence’s intermittent periods (SM).

## SIMULATED VS. REAL-WORLD NETWORKS

As an attempt to understand the fundamental workings of the human brain, spiking dynamics of biological neural networks has attracted considerable scientific interest. One important observation is that these networks often operate in or around a critical regime. In this case, spike dynamics is characterized by firing activity clustered in avalanches, with power-law size and waiting time distributions [6, 7]. Due to the diverging correlation length, network details are marginalized and a precise characterization of their behavior can be given and extended, by universality class-dependent scaling properties, into neighboring regimes [1]. Computational models proposed to capture this leave generally open to what extent they still faithfully reflect the properties of the real-world example beyond the critical regime.

Here, we test two widely used models (details see SM) of neural network simulation that both contain an avalanche critical regime. Rulkov neurons [24] can reproduce all salient experimentally observed spike patterns [25] and even neurobiological details such as phase response curves of biological neurons [26]. To model neuronal cultures, we consider networks of Rulkov neurons of size, composition and connectivity typical for cortical columns [8, 27]. Spike activity is transmitted along network edges by time-dependent synaptic currents, where their strength is scaled uniformly by a coupling strength parameter *W*. Individual Rulkov neurons are tuned to subcritical parameters, so that in absence of spontaneous or external input, firing activity dies out. Maturation of neural networks can be represented by an increase of the overall connectivity *W*, upon which a neuronal spike can trigger spiking of ever more connected neighbors. The network development with *W* has been shown to follow a universal pathway (i.e., is independent from the internal network wiring), explained in terms of simple dynamical systems elements and their interaction [8]. From the *n* neurons of the microcolumnar network (where normally *n* = 128, occasionally *n* = 256), *m* = 64 neurons were randomly selected to provide the ‘electrode signal’ of the real-world example. We find that the excess entropy of the Rulkov network’s firing activity deviates only slightly from the random walk baseline *H*_RW_(*n*), irrespective of the coupling strength *W* (Fig. 3 b,c).

**FIG. 3.**
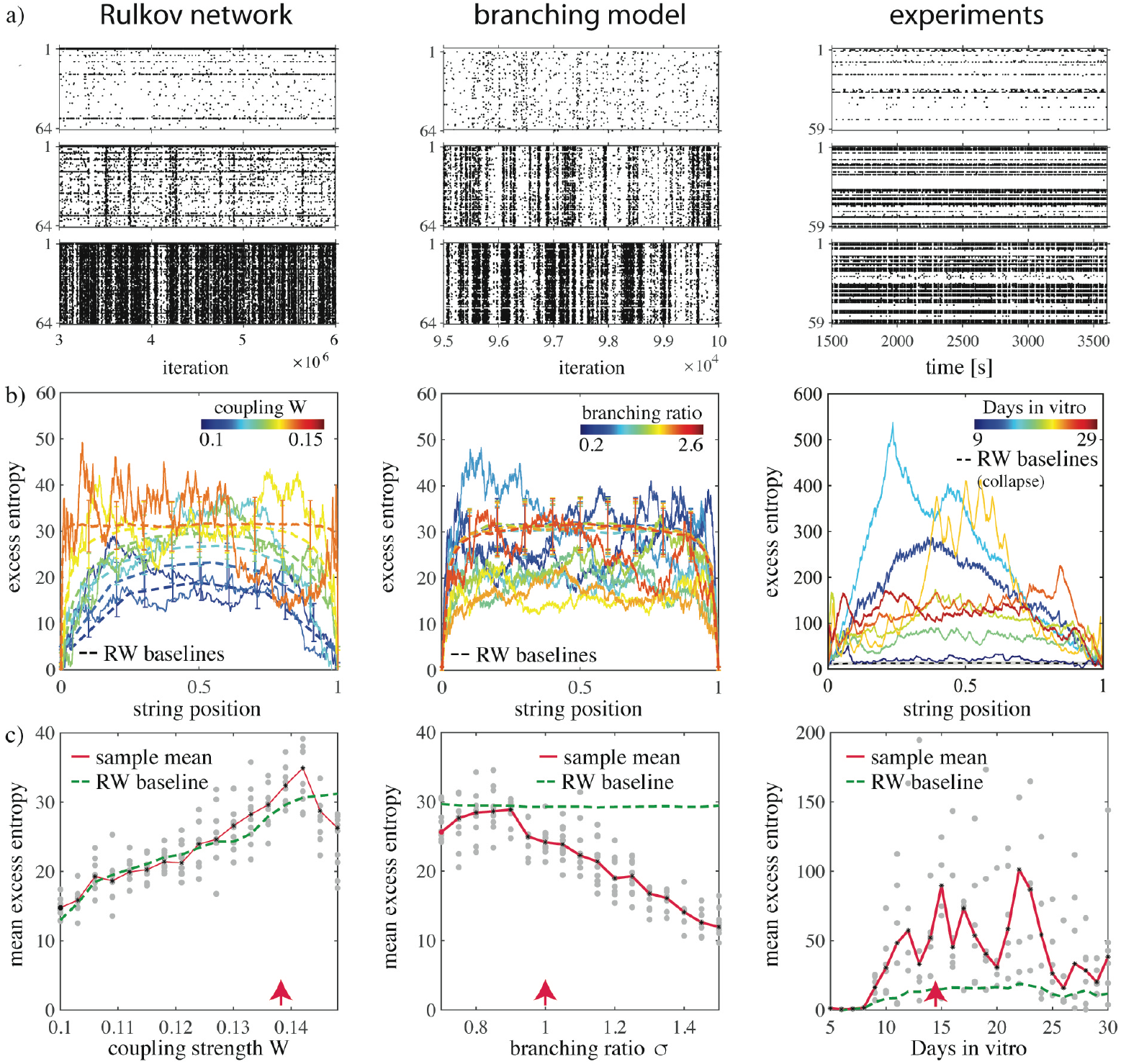
Simulated vs. real-world data: Excess entropies of both Rulkov and branching models (columns 1 and 2) are much lower than those of the biological example (column 3). a) Spike raster plots (*W* = 1.35, 1.39 (critical point), 1.5; *σ* = {0.7, 1 (critical point), 1.07}, cultures at 9, 13 (critical point), and 19 Days in vitro, each from top to bottom). b), c) While real-world entropies are way above, both modeling approaches yield, with excess entropies around or below the random walk baseline, much too regular stochastic dynamics. All cases contain avalanche critical points (approximate locations: red arrows).

A second model that may be expected to be even closer to an experimental MEA setting is the branching network model [28], a mesoscale ansatz where each electrode is represented by a node in a directed network. Each node *i* is a processing unit that can either be active (spiking) or inactive (silent). Activity is transmitted from active node *i* to node *j* according to probability *p*_*ij*_ chosen such that ∑_*j*_*p*_*ij*_ = *σ* is constant. Similarly to *W* in the previous example, a parameter *σ* (*branching ratio*) determines the expected number of newly active nodes triggered by the active node *i*; maturation of cultures is reflected as an increase of *σ*. For *σ* = 1, power-law distributed avalanche sizes emerge, whereas for *σ* ≷ 1, the network is in a supercritical and a subcritical phase, respectively. Exemplary activity of a network of 8 × 8 fully connected units is shown in Fig. 3a for *σ* = (0.7, 1, 1.07). For this model, the mean excess entropy 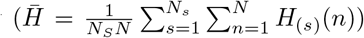 is not enhanced, but lies, after *σ* > 0.9, even substantially below the random walk baseline 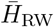. This demonstrates, compared to a corresponding simple random process, much more orderly firing dynamics.

We compared these simulation results with experimental data from developing neural cultures using our excess entropy method (a visual inspection of Fig. 3a provides little insight, due to the differing time-scales of the models). A total of six cultures of rat hippocampal neurons were grown and recorded on a 1mm^2^ 59-channel micro-electrode array chip (MEA), using a neural density of 800 cells / mm^2^. To minimize disturbance of the cultures, only a one-hour measurement was collected per day. A detailed description of the experimental setup and measurement protocol can be found in Ref. [29]. Our previous analysis of these recordings showed sparse or absent neural activity for DIV 5, leading to the exclusion of this data from the analysis. As the cultures mature, an increased firing activity emerges (Fig. 3a). We, moreover, previously showed that from around DIV 9 onwards, cultures start to exhibit power-law neural avalanche size and lifetime distributions in their firing patterns, where in Fig. 3 c, we have marked only the most prominent ‘late’ critical regime [29]. To apply the excess entropy measure, each electrode was associated with a unique state. Whenever electrode *j* registered activity above a fixed threshold, we considered the neural network to be in state *j*, resulting in a sequence **S** of transitions between the different states. Even though the information about the length of inactivity is not retained in this way, a high sampling rate ensured that the temporal order of electrode activity was correctly represented. *Surprisingly, our excess entropy analysis clearly demonstrates that both models deviate substantially from the experimental data, cf. Fig. 3c*.

To further investigate the saliency of this mismatch, we checked the consequences it has at the level of formal languages. While languages can be translated, a fundamental hierarchy classifies them from less complex to more complex, measured by their ease to capture complex relations among the language elements (Chomsky’s formal language hierarchy [10]). While the precise membership of a string to a formal language class is undecidable [13], we can circumvent this problem by a membership probability argument [17]. The simplest grammatical model for the putative generation of the experimental time series is a Chomsky hierarchy t-3 grammar. This model is equivalent to a random walk on the given set of symbols, with probabilities as observed in the respective experiments; strings of ‘noise’ of low complexity (complexity measured according to Ref. [16]), should therefore fit well into a random walk model. Comparing this model with the experimental excess entropies *H*(*n*) for six cell cultures shows that for all cultures at each DIV, the experimental values are considerably higher than those from the associated random walks (evaluated from a set of *N*_*sim*_ = 200 surrogate random walks). Moreover, the experimental values are also found to deviate strongly from those obtained from our two models of neural network simulations. The Average excess entropies 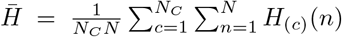 over each sequence length and all six cultures consolidate that the biological neuronal firing exhibits a strongly non-simply-random complex dynamics (Fig. 3c and Fig. 5 in our SM). From the developmental point of view, we generally find the smallest deviations from the random walk model for the initial (9 DIV) and late (25-30 DIV) stages, whereas strong deviations occur at the intermediate days, usually between 15-25 DIV. There, the behavior often switches between more random walk-like (for the shown example: 13 DIV and 16 DIV) and extremely complex characteristics (12 DIV and 15 DIV, Fig. 3b). In a previous study of the cell culture data [29] we found first fingerprints of avalanche criticality after 10 DIV, where our mean excess entropy shows a prominent increase, in agreement with what the logistic map example suggests (Fig. 2). The strong fluctuations of the excess entropy observed indicate that a simple t-3 grammar is unlikely to underlie the generation of the experimental data. Cultures of enriched medium are found to have even stronger deviations from the random walk model (SM).

To determine a potentially responsible mechanisms underlying the strong fluctuations, we scrutinize an experimental culture with strong variations of the excess entropy (DIV 22; yellow line in Fig. 3; other DIV and cultures exhibit parallel features). We find that the shape of the entropy separates into alternating regions, in which the entropy increases or decreases in a piecewise manner (see *h*(*t*) of DIV 22 from 1800 s to 2000 s in Fig. 4). Using time instead of string position as the abscissa, the spikes occur in distinct firing avalanches or bursts that engage only subsets of the culture. If we group the data according to their general trend towards increasing or decreasing entropy, the four regions distinguished by colors in Fig. 4 a emerge. To test these regions for similarities, we evaluated the activation probabilities for each electrode for each of the regions separately. Representing electrode activation during entropy increasing/decreasing regions by circles/crosses, respectively, we obtain very similar activation probabilities for entropy-increasing as well as entropy-decreasing regions (Fig. 4b). *This suggests that (increasing or decreasing) regions are based each one on one specific pattern of activity (with, for increasing or decreasing entropies, distinct ‘alphabets’) that are repeated across the respective displayed windows*. Similar albeith less striking activity patterns occur beyond the discussed observational window, where local maxima and minima of the entropy signal a change of the symbol probabilities.

**FIG. 4.**
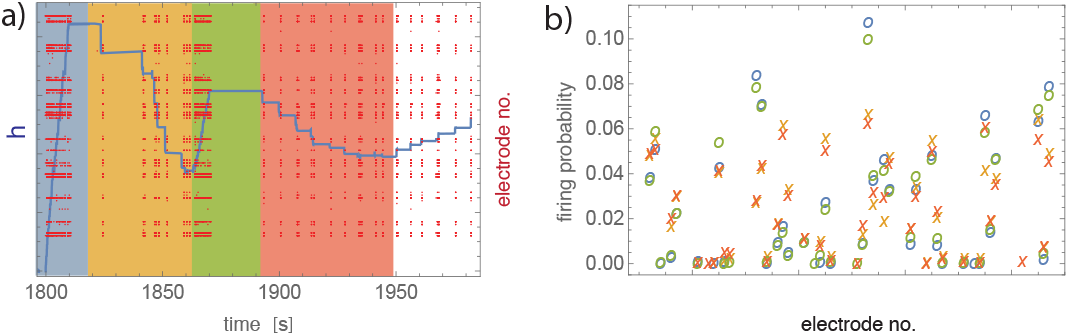
Zoom into 22 DIV data (*t* = 1800 s to 2000 s). a) Entropy *h* (blue, left axis) and corresponding electrode raster plot (red, right axis), where color shading indicates regions of increasing or decreasing entropy. b) Electrode activation probability for each of the four shaded regions of a), with symbol coloring inherited from a). Similar probabilities of circles coding for entropy-increasing (and similar of crosses in the case of entropy-decreasing) regions, suggest that only a single avalanche pattern is responsible for entropy increase (and for entropy decrease, respectively).

**FIG. 5.**
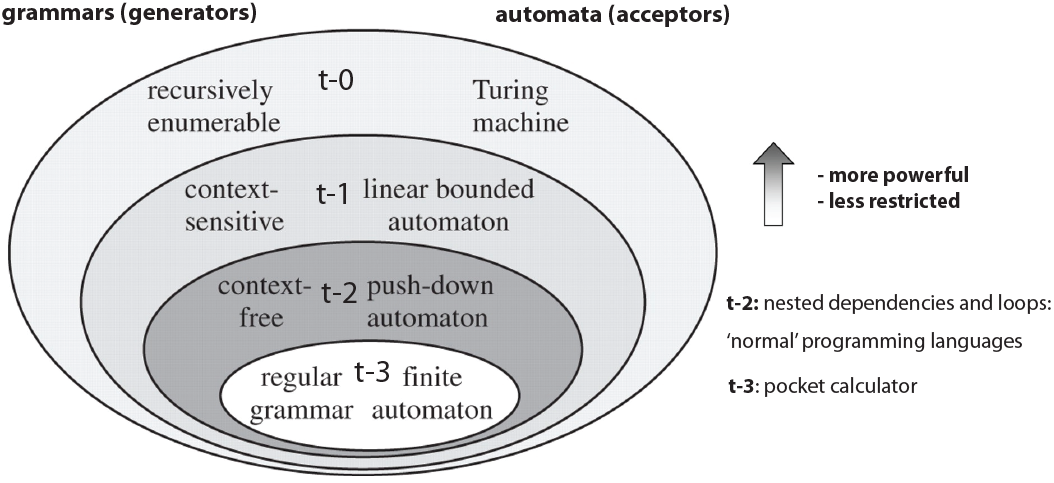
Chomsky language classes t-3–t-0 with corresponding properties. The inclusion property of the classes impedes a direct determination of the class that generated a string.

## CONCLUSION

We have presented a general tool that directly measures the distance between symbolic dynamics of simulations to those of the real-world example. By measuring how much simulation-generated symbol substring distributions deviate from distributions generated by the experimental example, also temporally variable patterns of transitions (that are generally difficult to extract from experimental systems) are included, which reaches beyond classical transition-matrix based methods. Strongly positive deviations indicate increased temporal clustering of subsets of symbols, whereas strongly negative deviations indicate that symbols occur in a too regular fashion. Such information exhibits weak points in the modeling and, as a consequence, can be used as a guideline for improving the modeling approach. As one application, our approach indicates that the Rulkov neuron modeling is much closer to the MEA experiment than the branching model (cf. excess entropies). After criticality at *W* ≃ 0.13, multiple modes of independent subnetworks emerge that, upon the increase of *W*, are bound by and then are released from sets that share synchronization in an intermittent-like manner [8], rather similar to what we observe in the biological example. The failure of this model to reach the experimental excess entropy levels suggests that its parameter *W* should be changed into a temporally variable quantity [29].

Excess entropy evaluation captures important finegrained signal properties that common approaches are insensitive to. With excess entropies consistently larger than in simulations, the analysis of a set of neural cultures reveals that the organization of spikes in developing neural cultures’ may be more ordered and more complex [16] than what simulations predict: Instead of a t-3 language class indicated by simulations, large positive deviations from the random walk baselines hint at a language compiler of stronger Chomsky class. This feature suggests that already neural cultures may be primed to deal with data that request for their efficient representation logical switches and loops that in the realm of artificial computation require stack-like constructs.

To render a distinction between dynamical systems finer, we can generalize the approach to order-*q* excess entropies, with *q* ∈ (−∞, +∞), according to

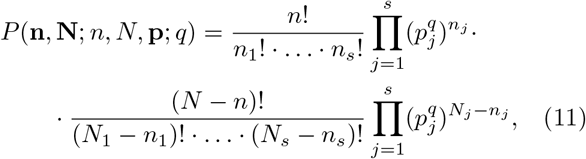

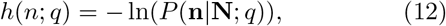

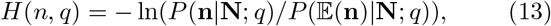

which offers a yet increased fine-grained analysis.

Based on our findings, we suggest our approach to be used to develop and validate more realistic simulations of neural networks, and generally of systems with similar characteristics. On the biological side, the challenge is to identify the detailed biological mechanisms responsible for the discrepancy between modeled and biological neural networks.

## Acknowledgments

The authors thank the group of Y. Nam, Kaist, Daejeon, Korea, for the neural cultures data. We also acknowledge the Swiss National Science Foundation SNF (grant IZKS2 162190) that made this work possible.

## Author contributions

N.S and R.L.S. share equal first authorship. R.S. provided the idea of the method. A.K. provided the processing of the biological data. R.L.S. developed the idea, contributed models and analyzed the data. N.S. provided the mathematical background and formulation. All authors wrote the paper.

## Correspondence and requests

Ruedi Stoop.

## SUPPLEMENTARY MATERIALS

### t-3 formal language class failure of cultures

The Chomsky hierarchy [10] defines classes of computational models in terms of formal languages called (type) t-3, t-2, t-1, t-0, each one a subclass of the next, increasing - along the presented order - in complexity and power, cf. Fig 5. Of central interest to us are class t-3 (representing the power a simple pocket calculator) and class t-2 (that represents the power of essentially all modern programming languages by offering structures of switches and loops).

To judge symbolic data from this angle, we first check whether the data is statistically satisfactorily represented by a random walk on the symbols, using the given symbol probabilities. If this holds, the data has t-3 class membership [10]. If not, the deviations from the random walks will look as in Fig. 3b, right panel. In the latter case, the data can be treated as follows: At the point of maximum *h*(*x*), the string is split and the resulting partial strings *ω*_1_ and *ω*_2_ are modelled separately. Since strings of the form *ω* = *ω*_1_*ω*_2_ require at least a t-2 grammar [17], experiments that are reproduced in a probabilistic sense (via comparison to surrogate data) by two partial random walks, are associated with a t-2 grammar (yet unclassified data sets are treated similarly by splitting non-matching substrings at the position of maximum entropy; they are associated with t-1 class membership [17]). Our experimental data falls clearly in a class stronger than t-3 (cf. Fig. 6).

**FIG. 6.**
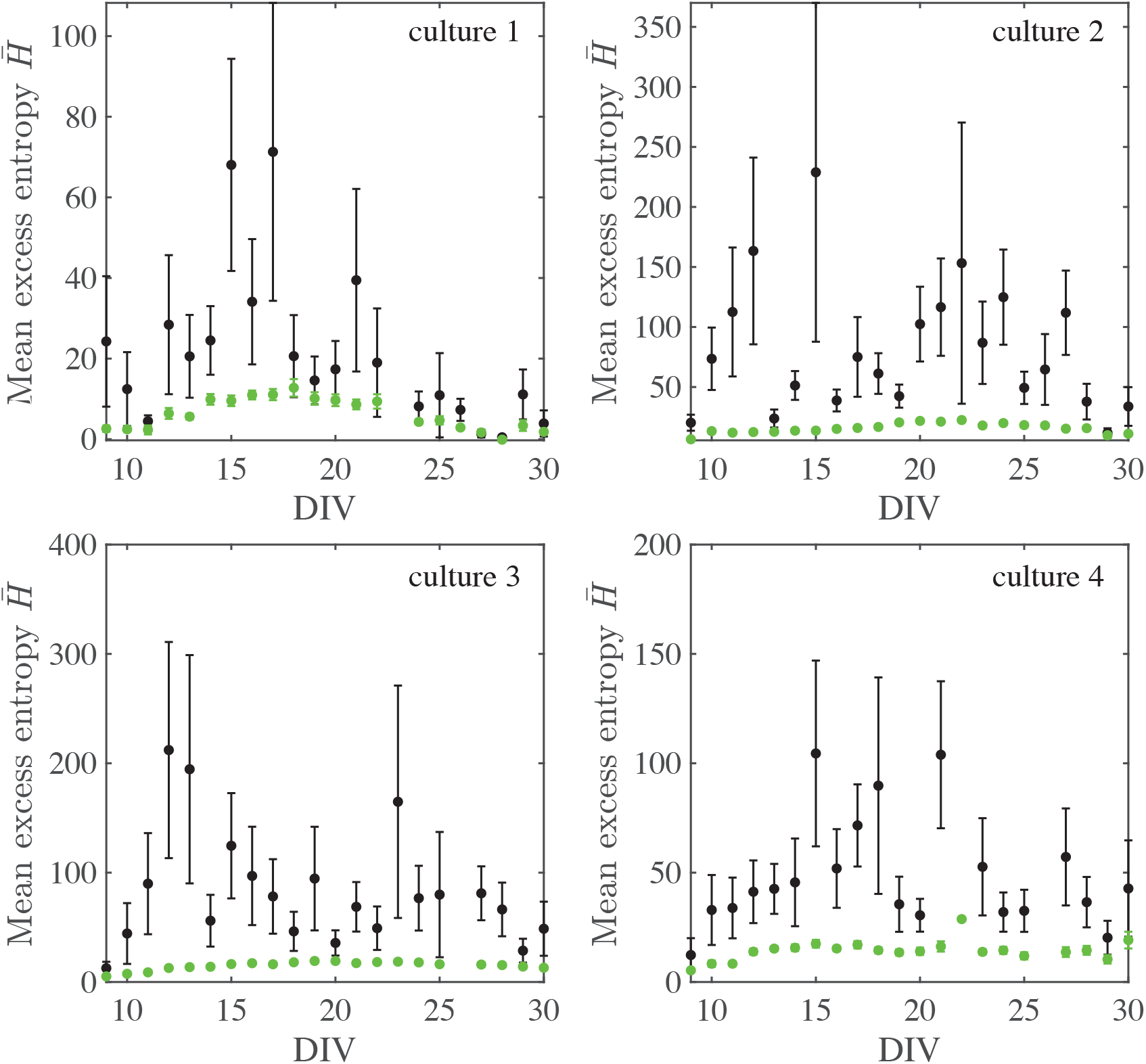
Excess entropies 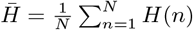 averaged over the length of the data string for four more neural culture electrode recordings (mean: black) and corresponding random walks (mean: green). Vertical bars denote one standard deviation from the mean along the string; intervals of random walks are found to collapse.

### Flip ratio of dynamical systems

The flip ratio is defined as

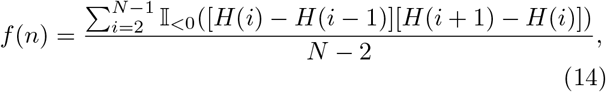

where 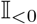 is the indicator function with value 1 if its argument is negative and zero otherwise. Since *f*(*n*) is a simple measure of tangent-tangent correlation, for a random walk, we expect 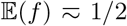. For the intermittent family, we find for *z* = 1 a value smaller than 1 (cf. Fig. 7), indicating the large degree of correlations that are present in the Bernoulli map (making it actually a bad random generator).

**FIG. 7.**
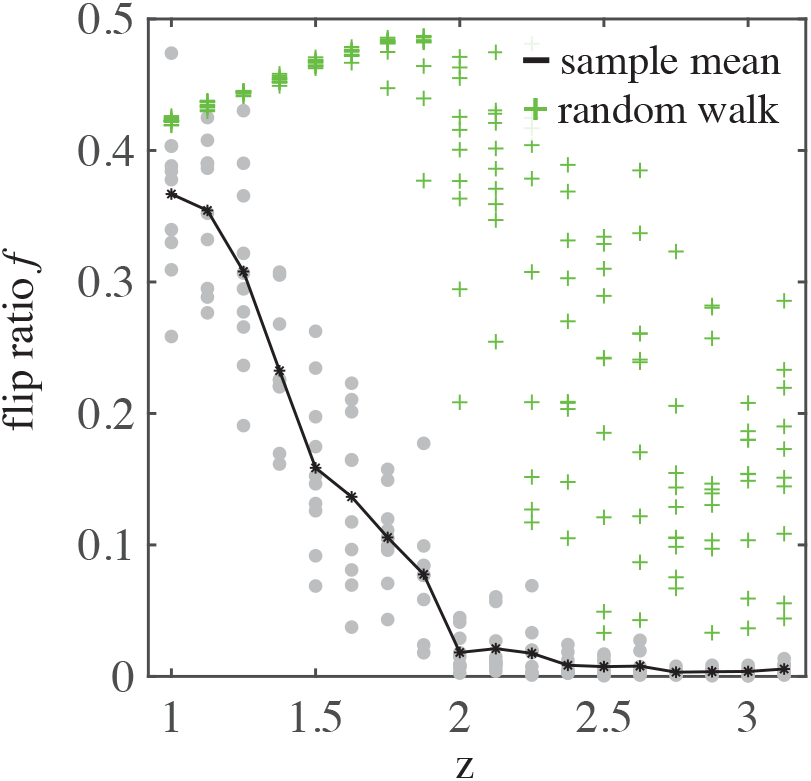
Flip ratios evaluated across blocks of size *L* = 4000 for the intermittent map family compared to a corresponding random walk of the length, in dependence of the exponent *z* of the map at the fixed point. Since the dynamics remain in the vicinity of the fixed point, for *z* ≥ 2, the flip ratio approaches zero.

### Asymptotics of 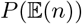

We consider a string of length *N* with *s* symbols occurring *N*_*s*_ times with probabilities **p** = **N**/*N*. Fixing a position *n* = *N*/2 in the middle of the string for simplicity, Eq. 4 is

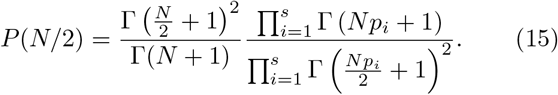

For *N* large, the Gamma functions can be expanded as

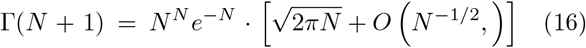

from which we obtain for the first fraction in Eq. 15

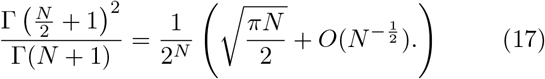

Using ∑_*s*_*p*_*s*_ = 1 and some algebraic manipulations, we find for the second fraction in Eq. 15

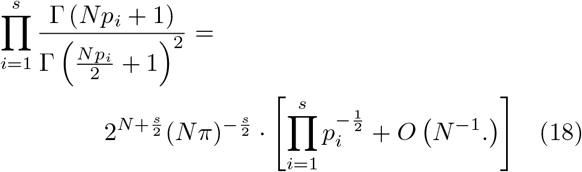

Combining both fractions, we obtain after collecting and combining terms of equal order

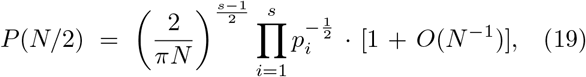

we immediately see that for *N* → ∞, 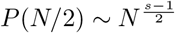.

### Lower bound

The asymptotics derived in the last section were based on symbolic strings where each symbol occurs in the middle of the string precisely as often as expected from probability. Such strings also provide an approximate lower bound for *h*(*N*/2) or, equivalently, the maximum of *P*(**n**|**N**). Since we only have to consider permutations of substrings of given **N**, we consider the walk-through probability *P*(**n**, **N**, *n*, *N*, **p**), which we express, as before, by means of the Γ-function. For any string **S** of length *N* with *s* symbols and probabilities *p*_*i*_, *i* = 1,…, *s*, we have

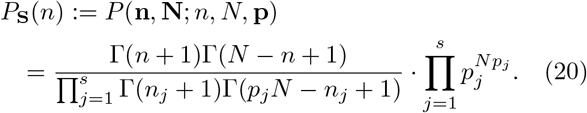

We compare this with the string **Ŝ** based on the same symbols and symbol probabilities, but where at each position *n* symbol *j* occurs exactly 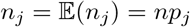 times. The corresponding probability of **Ŝ** is

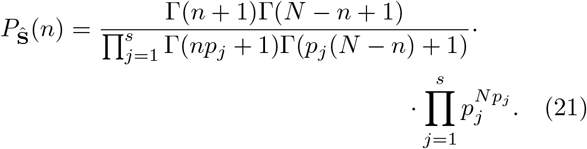

We first show that *P*_**Ŝ**_(*n*) ≥ *P*_**S**_(*n*) for all strings **S** having identical symbols and symbol probabilities as **Ŝ**. We start by considering a string **S** identical to **Ŝ** except that at position *i*, symbol *s*_1_ is interchanged with symbol *s*_2_ in **S**. In order for both strings to have the same number of symbol occurrences in total, this implies that at position *j* symbol *s*_1_ appears in **S** and *s*_2_ in **Ŝ**. Without loss of generality, we assume *i* < *j*. Thus, *P*_**S**_(*n*) = *P*_**Ŝ**_(*n*) for *n* < *i* and *n* ≥ *j*. For *i* ≤ *n* < *j*, the ratio of walkthrough probabilities at position thus is

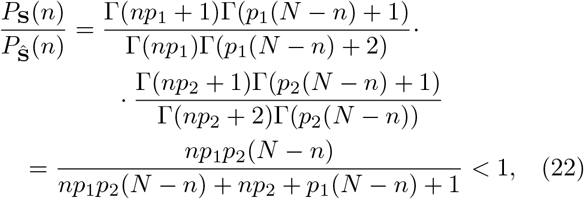

and therefore *P*_**S**_(*n*) < *P*_**Ŝ**_(*n*) for *i* ≤ *n* < *j*: Any permutation of two symbols results in a lower or equal walkthrough probabilities.

We now consider a string **T** differing from **S** by an additional symbol permutation *s*_3_ ↔ *s*_4_ at positions *k* and *l*, *k* < *l*. If the intervals [*k*, *l*] and [*i*, *j*] do not overlap, *P*_**T**_(*n*) < *P*_**Ŝ**_(*n*) for *n* ∈ [*i*, *j*] ⋃ [*k*, *l*] according to Eq. 22, with equality otherwise. We finally consider overlapping intervals such that *k* > *i*. For *n* < *k* and *n* ≥ min(*j*, *l*), it again follows from Eq. 22 that *P*_**T**_(*n*) < *P*_**Ŝ**_(*n*). In the overlap region *k* ≤ *n* < min(*j*, *l*), we find

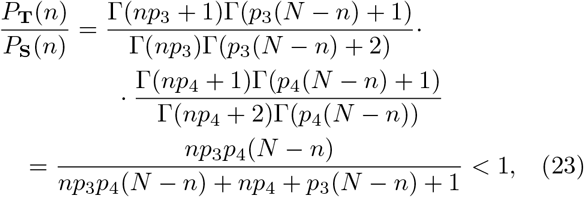

and therefore,

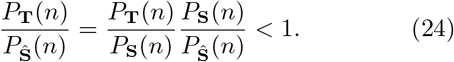

Since any permutation **Ŝ** of the string **S** can be expressed by successive pairwise permutations, *P*_**S**_(*n*)=*P*_**Ŝ**_(*n*) ≤ 1 for all strings **S**. It is important to note that we have shown in the above derivation that among the candidate occurrences *p*_*s*_*n* ± [0, 1, 2,…], the highest walkthrough probability is attained for occurrence *p*_*s*_*n*, which is generally not integer. Conversely, it is possible and numerically verifiable that an integer occurrence number *n*_*s*_(*n*) in the interval [*p*_*s*_(*n* − 1), *p*_*s*_(*i* + 1)] provides a slightly higher walk-through probability *P*. Since symbolic strings always have integer symbol occurrences, *P*_**Ŝ**_(*n*) can be an approximate bound only: Any string exceeding the bound at position *n* will have symbol occurrences *n*_*s*_(*n*) that differ from the bound occurrences *np*_*s*_ only by rounding up or down to the nearest integer. Due to the strong nonlinearity of the Gamma function, rounding to the nearest integer for several symbols can lead to small, but noticeable deviations from the bound.

### Excess entropy 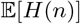 of random walks

For random walks, we have (Eq. 7)

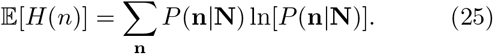

Due to the nonlinearity of the logarithm, 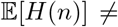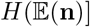. But even 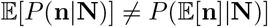. As specific example, consider position *n* = *N*/2 in the middle of a string of only two symbols *s*_1_ and *s*_2_. Assuming symbol probabilities *p* and *q* = 1 − *p* respectively, Eq. 11 is the product of two binomials

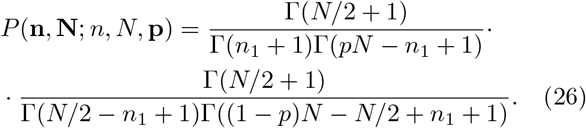

For long strings and probabilities *p* and *q* relatively similar, *n*_1_ is a random variable with approximately binomial distribution with *N*/2 trials and probability *p*. Under this assumption, the expectation value 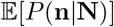 is

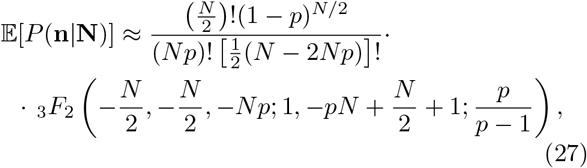

where _3_*F*_2_ is the generalized hypergeometric function with 3 and 2 parameters of type 1 and 2, respectively. This is obviously different from 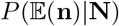. A numerical evaluation for *N* = 300, *p* = 1/3 is shown in Fig. 8 (blue).

**FIG. 8.**
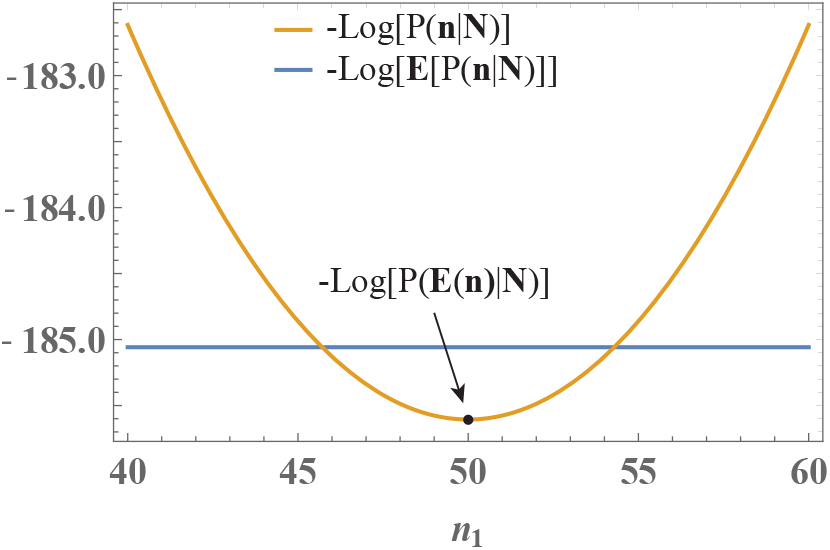
Comparison between logarithms of expected random walk probabilities 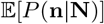 (blue) and *P*(**n**|**N**) in the middle of a sequences containing only 2 symbols with occurrences *n*_1_ and *n*_2_ = *N* − *n*_1_ (*N* = 300, *p*_1_ = *p* = 1/3). In the middle of the string, *P* is highest if symbol 1 is encountered exactly *pN*/2 = 50 times, which is considerably larger than the expectation value 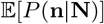.

### RULKOV MODEL

In our Rulkov neuron networks, each neuron is modeled by a two-dimensional Rulkov map [24]

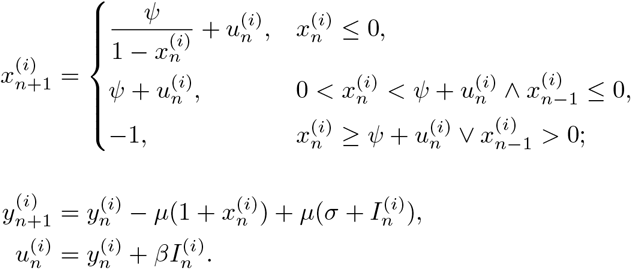

Here, *n* is the iteration step, 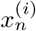 the membrane potential of the *i*’th neuron, and 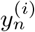 describes a regulatory sub-system able to turn firing on and off. *ψ*, *μ*, *β*, and *σ* are scalar parameters. Neuronal spike events are modeled by threshold variables

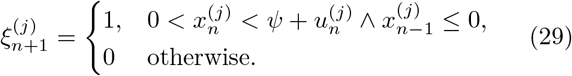

Lastly, 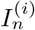 is the synaptic input to neuron *i* [25],

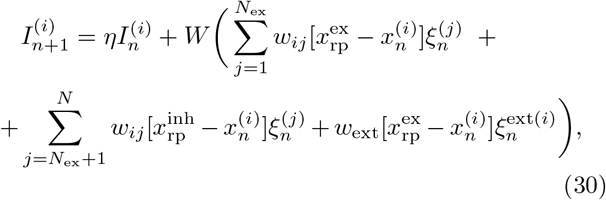

with decay rate *η* < 1, and reversal potentials 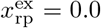 and 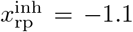 of excitatory and inhibitory synapses, respectively. A network is described by a connectivity matrix *w*_*ij*_, where *w*_*ij*_ = *w*_ex_ (*w*_*ij*_ = *w*_inh_) if neuron *i* is the terminal of an excitatory (inhibitory) synaptic connection from neuron *j*, and *w*_*ij*_ = 0 otherwise. Parameter *W* is a coupling strength, similar in function to the branching ratio in the branching model. Rulkov dynamics hosts a weak type facilitation and depression: facilitation by *I*_*n*_ renders Rulkov neurons more excitable (in the case of excitatory input) by elevating the *x*_*n*_ variable and by depressing the *y*_*n*_ variable.

### BRANCHING MODEL

We implemented the branching model as outlined in Ref. [28]. We considered simulations with *N* = 64 processing units, but our results remained essentially identical if we randomly choose 64 read-out units from a network of *N* = 1024 units. In accordance with Ref. [28], we choose a fully connected network of units, i.e. each unit is connected to all other *N* − 1 units. A randomly chosen but fixed probability *p*_*ij*_ of transmitting is assigned to each connection. If unit *i* is active, the branching parameter *σ*_*i*_ = ∑_*j*_*p*_*ij*_ determines the expected number of activated descendants. Probabilities are then rescaled such that the branching parameter is the same for all units. We also added spontaneous activity to the network by randomly activating any unit with probability *p* = 0.001.

